# A TetR-like transcriptional regulator in *Stenotrophomonas maltophilia* involved in fatty acid metabolism is controlled by quorum sensing signals

**DOI:** 10.1101/2022.08.31.506136

**Authors:** Xavier Coves, Pol Huedo, Marc Bravo, Òscar Conchillo-Solé, Andromeda-Celeste Gómez, Anna Esteve-Codina, Marc Dabad, Marta Gut, Xavier Daura, Daniel Yero, Isidre Gibert

## Abstract

*Stenotrophomonas maltophilia* is an environmental bacterium and it is also an emerging opportunistic multidrug-resistant pathogen. It uses the endogenous DSF quorum sensing (QS) system to coordinate population behaviors and to regulate virulence processes but can also respond to exogenous AHL signals produced by neighboring bacteria. Whole-transcriptome sequencing analyses were performed for *S. maltophilia* K279a in the exponential and stationary phases as well as in exponential cultures after treatment with exogenous DSF or AHLs. The results revealed that at the beginning of the stationary phase 1673 genes are differentially expressed. COG analysis showed that most of these genes were enriched for energetic metabolism processes and regulation of gene expression. After adding DSF or AHLs, 28 or 82 genes were found deregulated, respectively, 22 of which upregulated by both autoinducers. Interestingly, among these later genes, 14 were also upregulated in the stationary phase. Gene functions regulated by all conditions include lipid and amino acid metabolism, stress response and signal transduction, nitrogen and iron metabolism, and adaptation to microoxic conditions. Among the common top upregulated QS core genes, a putative TetR-like regulator (Smlt2053) was selected for functional characterization. This regulator has been shown to control a narrow regulon, including its own operon. It was found to sense long-chain fatty acids, including the QS signal DSF, and regulate a β-oxidation catabolic pathway. Overall, our findings provide clues on the role that the QS could have in *S. maltophilia* in the transition from the exponential to the stationary phase and bacterial fitness under high-density growth.

**IMPORTANCE:** The quorum sensing system in *Stenotrophomonas maltophilia*, in addition to coordinating the bacterial population, controls virulence-associated phenotypes, such as biofilm formation, motility, protease production, and antibiotic resistance mechanisms. Biofilm formation is frequently associated with the persistence and chronic nature of nosocomial infections. In addition, biofilms exhibit high resistance to antibiotics, making treatment of these infections extremely difficult. The importance of studying the metabolic and regulatory systems controlled by quorum sensing autoinducers will make it possible to discover new targets to control pathogenicity mechanisms in *S. maltophilia*.

## INTRODUCTION

To rapidly adapt to environmental changes, bacteria modulate complex social behaviors through physiological processes collectively known as quorum sensing (QS). QS is a type of cell-cell communication that allows individuals of the same species or different species to collaborate and compete through production, release, and response to extracellular signals known as autoinducers (AIs) (1). By sensing these AIs, bacteria detect and respond to cell density through gene regulation. As a result, QS enables bacterial populations to synchronize gene expression to coordinate biofilm formation, virulence and antimicrobial production and resistance, among others (2–4). Depending on the AI molecule there are different QS family variants that may be widely distributed among bacterial species or be exclusive to one taxonomic group (5). At the same time, many species possess and interpret more than one QS system, while others display only one type of AI signal.

In the ubiquitous Gram-negative bacterium *Stenotrophomonas maltophilia* the main QS system described so far is mediated by the fatty acid signal *cis*-11-methyl-2-dodecenoic acid, also called DSF (for <u>D</u>iffusible <u>S</u>ignal <u>F</u>actor). *S. maltophilia* has emerged over the last decades as an opportunistic pathogen, causing nosocomial infections in the immunocompromised population and it is commonly isolated from the lungs of cystic fibrosis patients (6, 7). One of the main concerns about the members of this species is that they tend to be intrinsically resistant to multiple antibiotics, with specific isolates showing evidence of adaptive resistance (8, 9). In addition, they have the ability to attach to abiotic and biotic surfaces and colonize medical devices and epithelial tissues forming biofilms (10), and are prone to acquiring virulence factors through horizontal gene transfer (6). The DSF-mediated QS system has been proven to control biofilm formation, motility and production of virulence and resistance factors in *S. maltophilia* (11–13), and a link has been established between DSF and susceptibility to some antibiotic classes in clinical strains (14).

The DSF-based QS system was first described in *Xanthomonas campestris pv. campestris* (*Xcc*) (15) and later reported in other opportunistic pathogens such as members of the *Burkholderia cepacia* complex. Signal molecules of the DSF family mediate intra- and inter-species signalling in these species through components encoded in the *rpf* cluster (<u>R</u>egulation of <u>P</u>athogenicity <u>F</u>actors). In *S. maltophilia* the *rpf* cluster consists of two contiguous but convergent operons: *rpfFB* and *rpfCG* (11). Unlike other species, the distinctive feature of the DSF system in *S. maltophilia* is the presence of two allelic *rpf* variants (namely *rpf*-1 and *rpf*-2) (11, 12). Strains with the *rpf-1* variant are more abundant and they produce detectable DSF levels under laboratory conditions, whereas strains with variant *rpf-2* do not. However under certain circumstances, for instance, when high concentrations of exogenous DSF are present in the microenvironment, strains of the *rpf-2* variant produce DSF endogenously (12). The genetic and biochemical mechanisms of DSF signaling, including DSF turnover, have been extensively studied in several members of the *Xanthomonadaceae* family (16–21). However, in *S. maltophilia* little is known about the regulation of this QS system.

Beyond DSF, the most common QS system in Gram-negative bacteria is based on N-acyl-homoserine-lactones (AHLs) (22). Despite AHL activity has been associated with members of the *Stenotrophomonas* genus (23, 24), *S. maltophilia* does not express the classical AHL synthases (25, 26), and to our knowledge, no canonical AHLs synthases have been described in this species. However, *S. maltophilia* can respond to AHL signals through its LuxR solo SmoR, which recognizes N-(3-Oxooctanoyl)-homoserine lactone (3OC8-HSL) and enhances bacterial motility (27). On the other hand, *S. maltophilia* AHL-degrading strains have been isolated from the microbiota of aquatic organisms (28, 29), which indicates that these organisms can detect AHL molecules in the environment before they are hydrolyzed. The possibility of externally induce AHL degradation, a quorum quenching (QQ) activity, is attractive because it offers the possibility of modulating virulence of some pathogens by interfering with the QS in highly competitive niches, such as the rhizosphere or the lungs of CF patients (30–32).

Owing to the limited knowledge on the effect of exogenous AI molecules in *S. maltophilia* and the potential therapeutic use their inactivation may have, the global expression profiling of *S. maltophilia* exposed to QS signals that are typical in its most common natural niches is studied. To start deciphering this response, transcriptomic analyses were performed by RNA-Seq, comparing the expression profiles in two cell density-based stressing conditions: (i) stationary-phase of growth and (ii) growth in the presence of QS signals. Subsequently, genes that could be relevant for the pathogenesis and persistence of these bacteria, as well as those that could regulate or be involved in signal turnover, were further investigated. A tetR-like regulator was found to sense fatty acids including DSF and drive this signal into fatty acid catabolism. This work provides a comprehensive global picture of how *S. maltophilia* responds to signals emitted by neighboring bacteria and the impact on virulence, bacterial fitness and the strategies used to overcome these stressing situations.

## RESULTS

### Changes in gene expression in stationary phase cells of *S. maltophilia*

QS systems play a crucial role in cells as population density increases and stationary phase approaches. Global gene expression was first monitored in the early-stationary phase (24 h, OD_600_=5.0) of *S. maltophilia* K279a (*rpfF*-1 variant) and compared with the mid/late-exponential phase (6 h, OD_600_=1.5) (Fig. 1A, time points b and c, respectively). *S. maltophilia* are considered fast-growing organisms and strains usually reach the stationary phase of growth in 20-24 hours with a high cell density (33, 34). Differential expression analysis of the RNA-Seq samples collected at both phases showed a total of 1673 differentially expressed genes (DEGs), representing 37% of total genomic content of *S. maltophilia* K279a. Of those genes, 908 were found significantly upregulated, whereas 765 were found downregulated (the full list of significant DEGs is provided in the data supplemental material). When all DEGs were analyzed in terms of their functional categories (Fig. 1B), main differences were observed in COG categories related with metabolism and regulation of gene expression. For instance, inorganic ion transport, secretion, and replication, recombination and repair were the most enriched categories in the upregulated dataset. On the contrary, the fraction of genes assigned to motility, energy production, translation and ribosomal structure and cell wall biogenesis represented the most enriched categories in the downregulated dataset.

**Figure 1.**
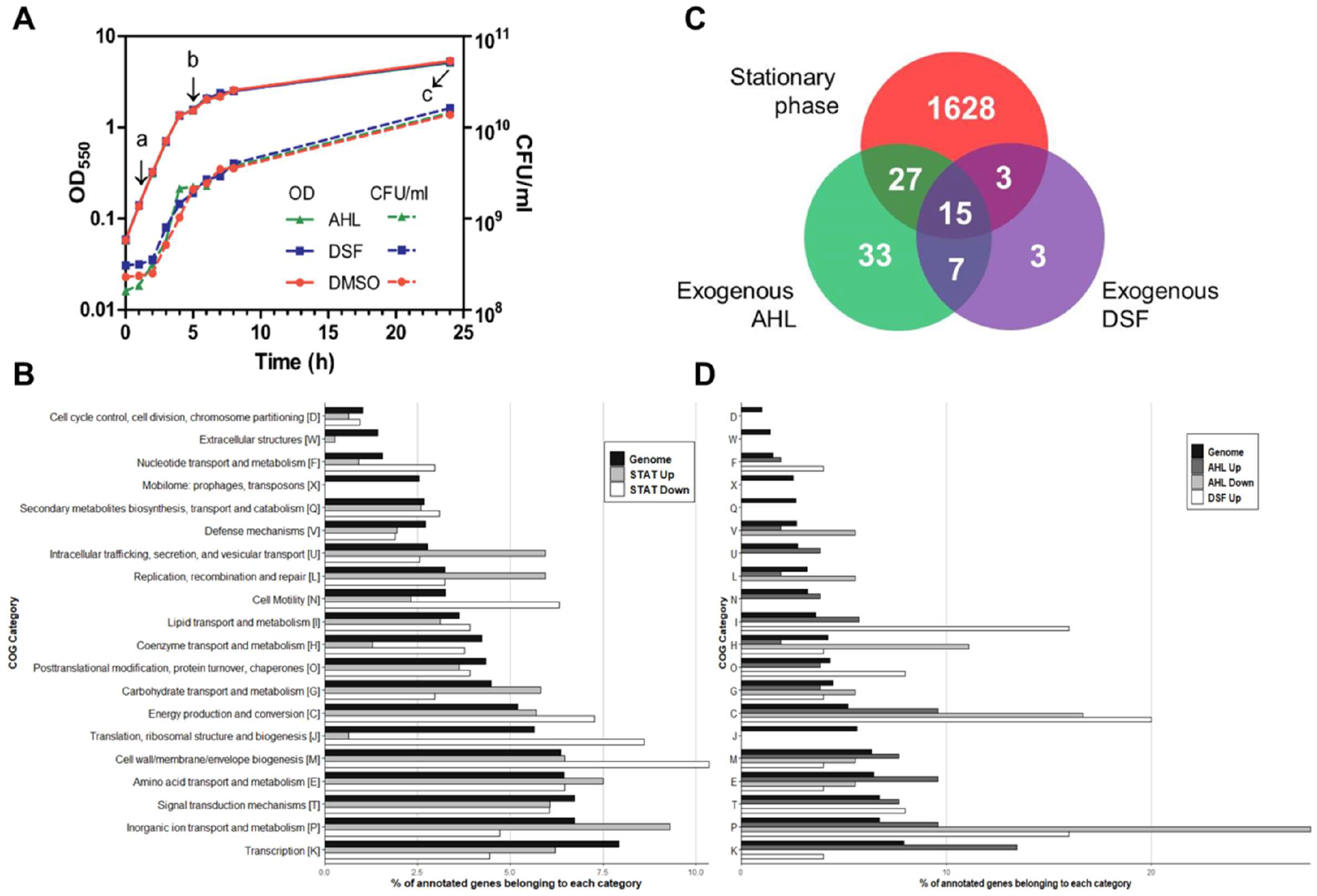
Experimental design and global analysis of differentially expressed genes (DEGs) under QS-inducing conditions. (A) Growth curves of the K279a strain in each condition, supplemented with AHLs, DSF or DMSO as vehicle control. Cultures were induced at time point (a), the start of the exponential phase and harvested at point (b), which corresponds to the mid/late-exponential phase of growth. A non-induced culture was also harvested at this point (b) to be used as the exponential phase representative (LOG) and to be compared with that of the early-stationary phase (STAT), harvested in (c). The growth curve of the non-induced culture was omitted for better visualization. (B and D) Functional classification in Clusters of Orthologous Groups (COG) of DEGs. COG categories are indicated in the y axis, while the x axis represents the percentage of genes matching each COG category at each condition, as well as in the whole genome (according to the IMG database). Genes with unknown function were not included in the graphs. Both the upregulated and downregulated genes are indicated in each condition with different colors. Growth-phase-specific differences in global transcriptome are shown in panel B. (C) Venn’s diagram showing overlap of DEGs in response to the three assayed conditions. (D) COG categories of DEGs after treatment with the autoinducer molecules. For the DSF condition only upregulated genes were detected. The full list of significant DEGs is provided in the data supplemental material.

Most of the top upregulated genes (>5 fold changes) belong to energetic metabolism, including genes involved in oxidation-reduction processes (e.g. *cta* operon encoding the cytochrome c oxidase and putative cytochrome *bd* subunit genes), DNA replication (putative *nrd* operon Smlt2839-Smlt2842), metabolite transport and secretion. The major extracellular protease StmPR1 (Smlt0686), the glycoside hydrolase Smlt2180 and the quaternary ammonium compounds efflux transporter Smlt2852 are among the most overexpressed genes in the early-stationary phase. A variety of transcriptional regulators and sigma factor family genes were also overexpressed at stationary phase, including the uncharacterized regulators Smlt2428, Smlt2711 and the TetR-family regulator Smlt2053. Particularly, the sigma-70 family RNA polymerase factor Smlt1750 (13.4-fold) could control stress response genes at the beginning of the stationary phase. Among the downregulated genes stand out members of the cell division cluster Smlt0747-Smlt0760 (*mur* and *fts* genes), the transcriptional repressor *lexA* and recombinase *recA*, two flagellum-associated operons, pilin-associated and chemotaxis genes, and the Ax21 family proteins Smlt0387 and Smlt0184.

### Effects of DSF and AHLs on gene expression

To gain insight into the transcriptomic response of *S. maltophilia* to a high-demand situation such as growing in the presence of QS signals, early exponential cultures were supplemented with naturally occurring concentrations of either DSF or an AHL combo, both dissolved in DMSO. Collection times were equal to those of the mid/late-exponential phase in the STAT vs LOG comparison (Fig 1A; b and c time points). Transcriptomic analysis revealed a total of 82 or 28 DEGs (p-adj<0.05, |fold-change| ≥ 2.0) after the treatment with AHLs or DSF, respectively, when compared to the DMSO control group (Fig. 1C). Upon AHL exposure, 58 genes were significantly upregulated and 24 genes downregulated, whereas DSF gave rise to only an inducing effect in the total number of DEGs. Top upregulated genes (>5 fold changes) in both conditions encode for the TetR-family regulator Smlt2053 and protein phenylalanine 4-monooxygenase.

COG analysis showed an enrichment in the categories of energy production and inorganic ion transport and metabolism for DEGs after treatment with both autoinducer molecules (Fig. 2D). DSF also upregulated genes related to lipid transport and metabolism, whereas AHLs induced amino acid and nitrogen metabolism. To develop a general picture of the biological processes affected by DSF and AHLs signals, GO enrichment analyses were performed on upregulated genes by both DSF and AHLs (Fig. S2 and S3). As with the COG analysis, oxidation-reduction and energetic processes were enriched in the common DEGs dataset.

**Figure 2.**
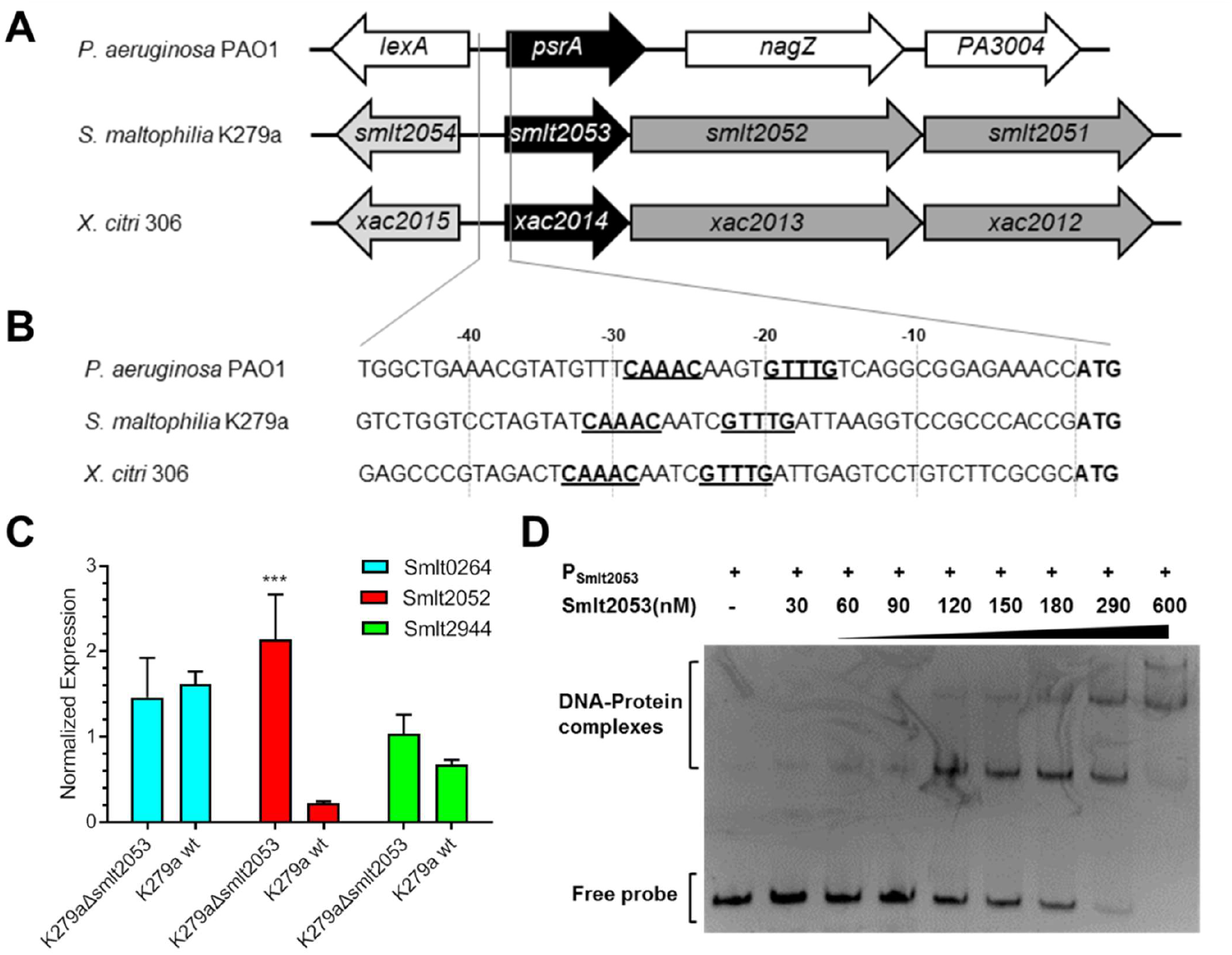
(A) Genomic context of orthologous TetR-like regulators in three Gammaproteobacteria. Gene names or locus tags are indicated in model genomes for strains PAO1 (NC_002516.2), K279a (NC_010943.1) and 306 (NC_003919.1). (B) DNA sequences of the operator regions of the three operons showing underlined the likely DNA recognition box. (C) Gene expression using qRT-PCR of Smlt0264, Smlt2052 and Smlt2944 in the *S. maltophilia* K279a Δ*smlt2053* (*tetR*) and WT normalized by the *rpoD* house-keeping gene. (D) Electrophoretic mobility shift assays for the promoter region of Smlt2053 (Psmlt2053) and the Smlt2053 protein. Molar protein concentrations are shown about the gel image and 50 ng of the DNA probe were used.

Most of the DEGs found in these conditions were also deregulated in the stationary phase of growth (Fig. 1D; 64% of DSF and 51% of AHL DEGs), which could indicate that the addition of exogenous QS autoinducers affects metabolic pathways in the same way as entering the stationary phase. For instance, the most downregulated genes after adding a cocktail of AHLs were also significantly repressed in the early-stationary phase. These genes belong to the *nar* operon (Smlt2769-Smlt2774), which encodes the respiratory nitrate reductase for nitrate respiration under oxygen-limiting conditions and includes the nitrate/nitrite transporter gene *narK*. On the contrary, an additional *narK* gene (Smlt2775), located immediately upstream to the *nar* operon, was found upregulated in the three conditions assayed. Smlt2775 is orthologous to *narK2* in *Pseudomonas aeruginosa*, which is exclusively required as a nitrate/nitrite transporter under denitrifying conditions (35). Contrary to *P. aeruginosa*, our results indicate that the transcription of *narK2* in *S. maltophilia* could be independent of the operon *narGHJIK1*.

Remarkably, the expression of 22 functional genes was induced by both QS signals, representing 79% of DSF and 38% of AHL DEGs (Fig. 1C and Table 1). The fold-regulation of four common induced genes (Smlt0077, Smlt0264, Smlt2053 and Smlt2944) was validated by qRT-PCR and the results closely mirrored the values obtained by RNA-seq (Fig. S1). In this data set, nine of the overexpressed genes are part of four operons (Table 1). One of these transcriptional units is a putative β-oxidation operon in the K279a genome (Smlt0264-Smlt0268), and the other is directly involved in iron storage (Smlt3600-Smlt3601). Interestingly, 14 out of these 22 genes were also found upregulated in the early-stationary phase (Table 1), including the TetR-family regulator Smlt2053 and the glycoside hydrolase Smlt2180.

**Table 1.**
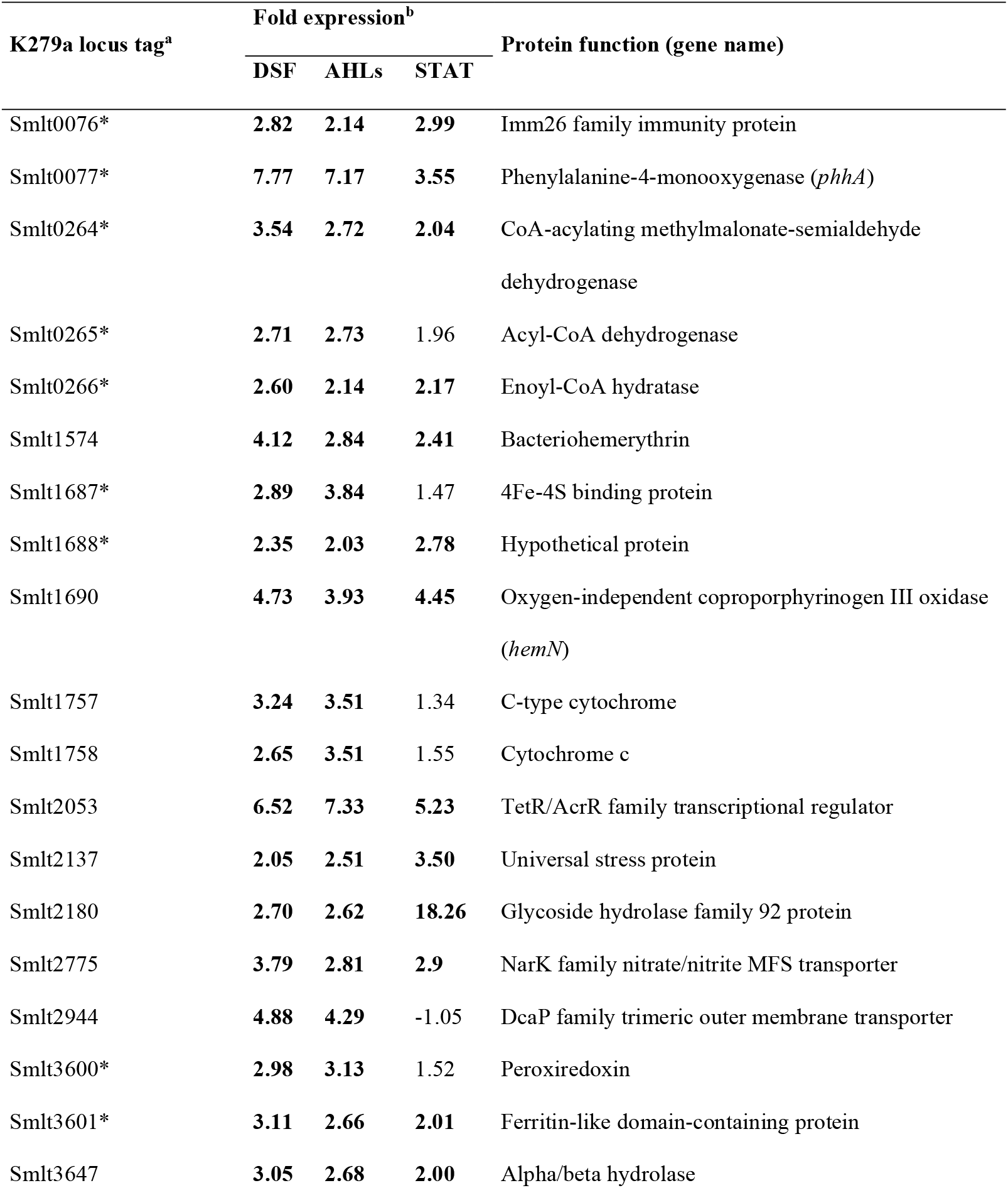
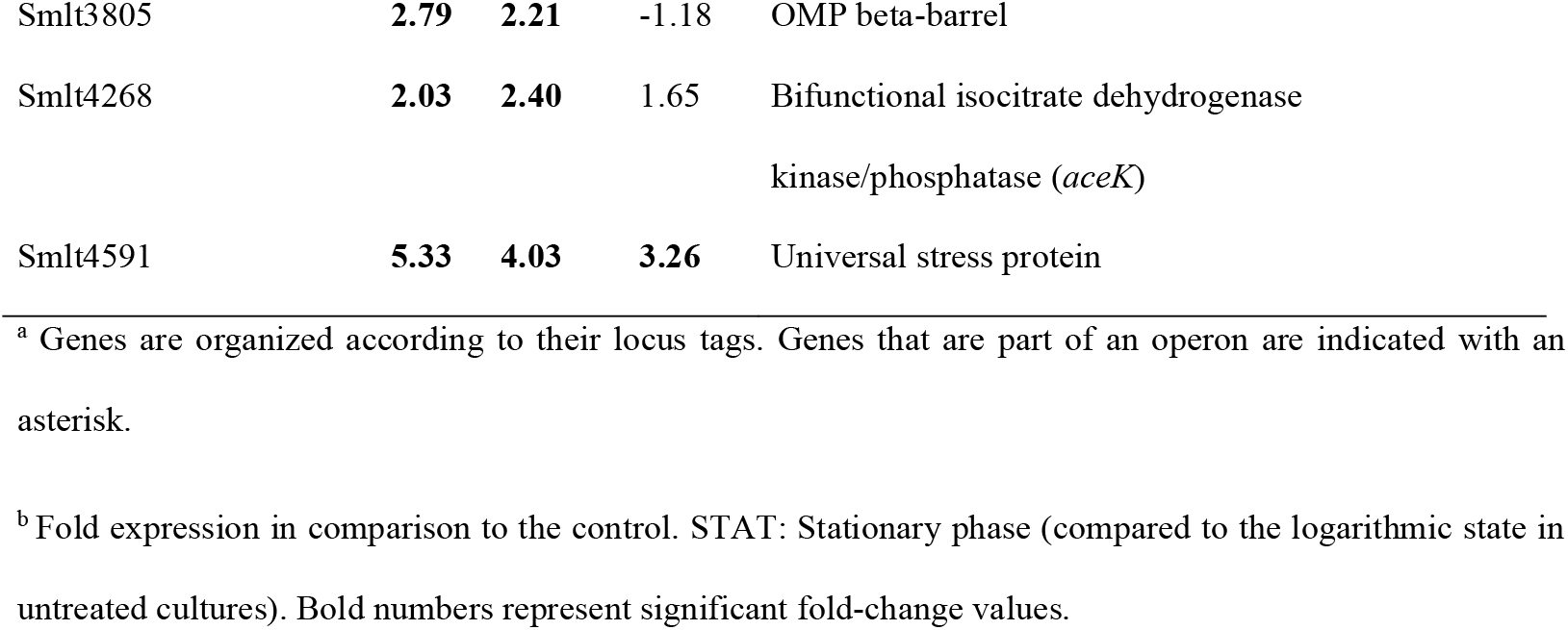
Common genes induced by both AHLs and DSF signals, compared to the expression profile during the early-stationary phase of growth.

### Smlt2053 is a transcriptional regulator that autoregulates its own expression

Among the common upregulated QS core genes (i.e., the genes deregulated in all conditions related to QS activation or sensing), we focused on the regulatory gene Smlt2053, as it was the first and the second most upregulated gene in the AHLs and DSF setups, respectively. In addition, this gene was also upregulated in the stationary phase of growth, with a fold increase of 5.23 (Table 1). It is annotated as a putative TetR/AcrR regulator; therefore, we initially hypothesized that it could act as a cytoplasmic regulator of the QS cascade or as a regulator of the quorum quenching pathway.

We initially performed sequence comparisons to identify homologues among related bacteria and obtain clues on its biological function. The closest regulatory protein with known function was identified in *Xanthomonas citri* (Xac2014) sharing 89% amino acid identity. Besides *Xanthomadaceae*, Smlt2053 also shares 39% identity with PsrA (PA3006) from *P. aeruginosa* PAO1. While *psrA* is organized as a single gene in the *P. aeruginosa* genome, genomic organization of Xac2014 and Smlt2053 is identical, with both genes conforming an operon encoding for an additional enoyl-CoA hydratase (Xac2013/Smlt2052) and acetyl-CoA C-acyltransferase (Xac2012/Smlt2051) (Fig. 2A), variants of the two enzymes of fatty acid catabolism FadB1 and FadA respectively (36).

The homology found between the three transcription factors is mainly restricted to the N-terminus, which corresponds to the DNA binding domain (37). In addition, the DNA recognition sequence of PsrA in *P. aeruginosa* has been identified as G/CAAAC(N_2-4_)GTTTG/C (Fig. 2B) (38). Notably, both Smlt2053 and Xac2014 present an identical DNA recognition sequence in their operator region and their position respect to the +1 is very similar. The similar N-terminus sequence of the three proteins as well as the identical DNA recognition sites found in their operator sequences suggests that the three transcription factors bind to their operator sequences repressing their own expression, as reported for PsrA (38–40).

We investigated the autoregulatory activity of Smlt2053 as well as the role that may exert on other QS-induced genes using different approaches. Firstly, a series of qRT-PCR were conducted in a *S. maltophilia* K279a mutant for gene Smlt2053 (K279aΔ*smlt2053*) compared with the WT strain. For mutant construction a markerless gene deletion was obtained without affecting its own promoter and thus avoiding the polar effect on genes Smlt2052 and Smlt2051 in the same operon. Three genes were selected for this analysis: Smlt2052, forming an operon with Smlt2053; and Smlt0264 and Smlt2944 because both were also found among the top 10 DEGs in the presence of the DSF and AHL signals. The absence of Smlt2053 resulted in a significantly increased expression of the adjacent gene Smlt2052 compared to the wild-type background, strongly suggesting its auto-repression activity and thus validating the operon organization of Smlt2053-Smlt2051 (Fig. 2C). On the other hand, no significant differences in expression were observed for genes Smlt0264 and Smlt2944, suggesting that their regulation would not be controlled by Smlt2053. To confirm the autoregulation of Smlt2053 we tested the direct interaction between protein Smlt2053 and its own promoter region *in vitro* using EMSA (Fig. 2D). The EMSA results showed that protein Smlt2053 specifically binds to a DNA fragment containing the promoter region, therefore indicating the binding of the regulator protein to its own promoter. The experiment revealed that as the concentration of the protein increases, an additional electrophoretically distinct protein–DNA complex with high affinity is formed. These findings suggest a different genomic target for the transcriptional regulator in the promoter P_Smlt2053_. Careful examination of the promoter region revealed an additional inverted repeat which resembles the consensus motif G/CAAAC(N_2-4_)GTTTG/C (Fig. S4).

To further investigate the role of Smlt2053 in QS and global regulation in *S. maltophilia*, additional RNA-seq experiments were performed on exponential cultures of the mutant K279aΔ*smlt2053* and the isogenic parent strain K279a and compared. Deletion of Smlt2053 affected the expression of only seven genes (p-adj < 0.05, |fold-change| ≥ 2.0) (Table 2). Five genes were downregulated in the mutant strain while the adjacent genes Smlt2052 and Smlt2051 were the only genes significantly upregulated, hence confirming the autorepression activity observed by qRT-PCR for this operon. Notably, the Bacteriohemerythrin gene Smlt1574, downregulated in the mutant Δ*smlt2053*, was found upregulated in the three previously assayed conditions (Table 1). The genes that were found more repressed in the mutant all belong to an operon (Smlt1443-Smlt1445) that also appears to be regulated by a different TetR-type regulator. An inverted repeat sequence similar to that found in the promoter of Smlt2053 was not found in any of the promoter regions of these DEGs in the mutant Δ*smlt2053*.

**Table 2.** DEGs in the mutant K279aΔ*smlt2053* compared to the WT variant.

| K279a locus tag | Fold-change | Protein function (gene name) |
| --- | --- | --- |
| Smlt1443 | -13.28 | HlyD family secretion protein |
| Smlt1444 | -11.50 | MFS transporter |
| Smlt1445 | -9.10 | TetR/AcrR family transcriptional<br>regulator |
| Smlt0598 | -2.36 | Asparagine synthase (*asnB*) |
| Smlt1574 | -2.06 | Bacteriohemerythrin |
| Smlt2051 | 6.02 | Acetyl-CoA C-acyltransferase |
| Smlt2052 | 6.35 | 3-hydroxyacyl-CoA<br>dehydrogenase/enoyl-CoA hydratase |

### Fatty acids including DSF affect the binding of Smlt2053 to its promoter

Since Smlt2053 is the ortholog to *P. aeruginosa* PsrA, a known TetR regulator of the β-oxidation pathway that binds fatty acids (41), we hypothesized that Smlt2053 would behave in a similar way. We use EMSA to monitor the effect of short, medium and long-chain fatty acids on DNA-Smlt2053 binding (Fig. 3). The presence in the mixture of short (C4) and medium (C8) length fatty acids did not affect the binding of the Smlt2053 protein to the probe containing its own regulatory region (data not shown). However, long chain fatty acids affected this binding, according to the EMSA results. When long chain fatty acids containing 12 or more carbons in its backbone were included in the binding reaction, with almost all the DNA probe P_*smlt2053*_ bound to the Smlt2053 protein, an increasing amount of free probe was observed correlating with the increasing amounts of each fatty acid (Figure 3). This confirms the ability of Smlt2053 to detect and bind long chain fatty acids, an initial step of the catabolic pathway. In addition, 13-methyltetradecanoic acid (iso-C_15:0_), the most abundant fatty acid in *S. maltophilia* which stimulates DSF synthesis (12), also reverted the Smlt2053-DNA binding (Figure 3).

**Figure 3.**
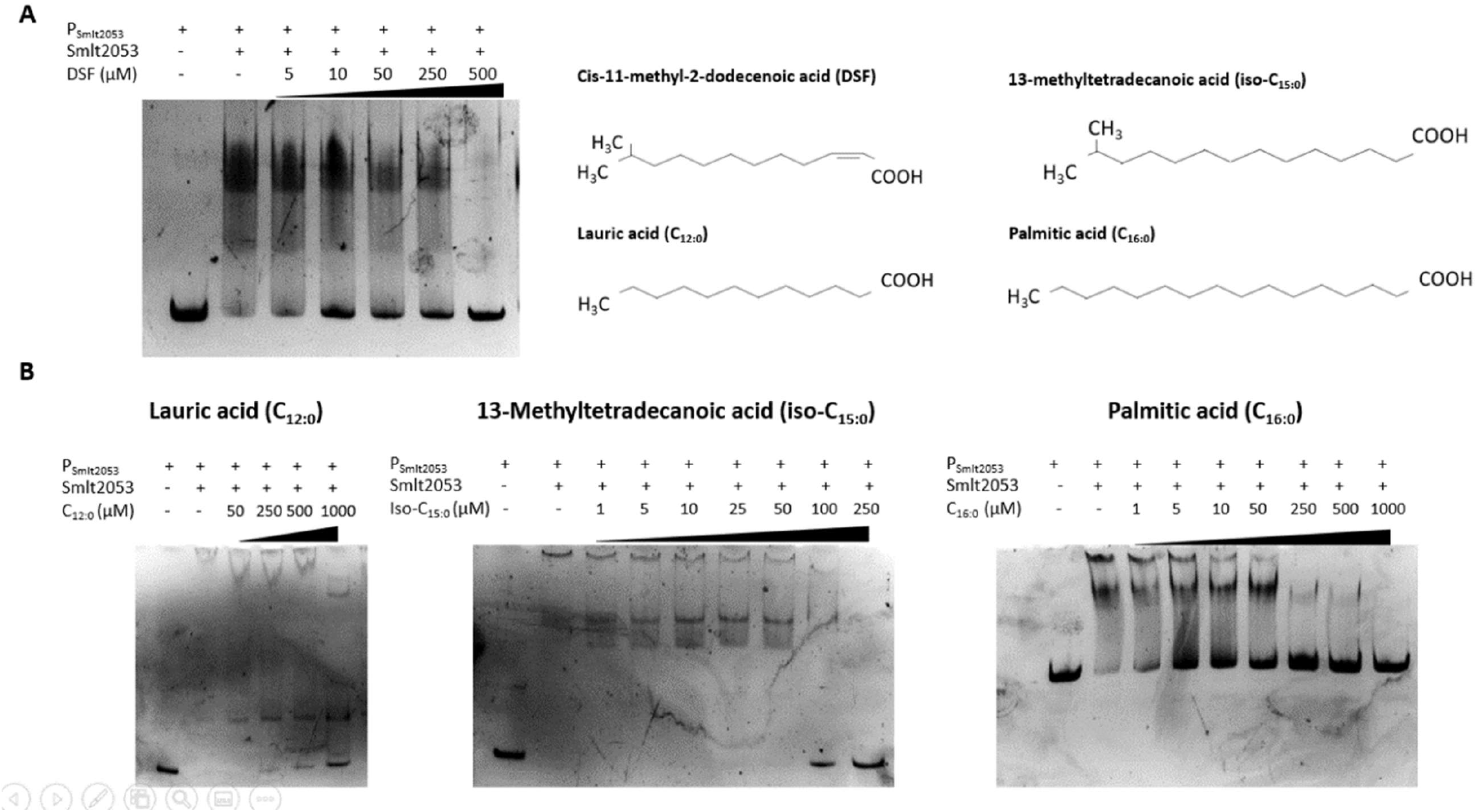
EMSA of Smlt2053-his tag protein with the DNA probe P_smlt2053_ in the presence of exogenous fatty acids. After 30 min of protein-DNA incubation, increasing amounts of each fatty acid were added to the reaction mixture for an additional 30 min. **(A)** DSF at concentrations ranging from 5 to 500 µM was added to the reaction. Lane 1 contained neither protein nor DSF and was used as a probe control. Lane 2 included protein and the same volume of DMSO (solvent) used at the highest concentration, as a vehicle control. In the right panel, the structure of each fatty acid used in this figure is indicated, to highlight similarities between them. **(B)** EMSAs in the presence of different fatty acids: lauric acid (C_12:0_), left panel; 13-methyltetradecanoic acid (iso-C_15:0_), center panel; and palmitic acid (C_16:0_), right panel. Lane 1 in all three experiments included only the DNA probe (P_smlt2053_). Lanes 2 included the protein and the vehicle used to dissolve each fatty acid, as follows: water containing 1% Brij-58 was used for C12:0 and iso-C15:0, whereas 10% methanol was used for C16:0.

Considering the lipid nature of DSF and its connection with iso-C_15:0_, we next examined if DSF could also act as a ligand for Smlt2053. Increasing amounts of DSF disrupted the shift observed between Smlt2053 and its own promoter region P_*smlt2053*_, indicating that Smlt2053 responds to the main QS signal of *S. maltophilia* (Figure 3). These results correlate to those obtained in the RNA-Seq experiments, where Smlt2053 was upregulated in the presence of DSF.

To study the effects that the operon regulated by the TetR-type transcription factor Smlt2053 may have on fatty acid metabolism, mutant strains for this protein regulator (K279aΔ*smlt2053*) and for all the components encoded by the operon Smlt2053-2051 (K279aΔ*smlt2053-2051*), which includes *fadB*/*A* gene variants, were evaluated. None of the mutants showed significant differences in terms of DSF production (data not shown). Neither the deletion of the components of this β-oxidation operon seem to alter the global fatty acid metabolism and regulation in *S. maltophilia* as determined by the fatty acid profile of K279a WT and mutants K279aΔ*smlt2053* and K279aΔ*smlt2053-2051* (Table S2). Fatty acid profiles with predominance of the 13-methyl-tetradecanoic acid (*iso*-15:0) are characteristic of the genus *Stenotrophomonas*, and no significant differences were observed in the abundance of all the molecules detected among the three strains analyzed. These results could also indicate that the enzymes of the operon Smlt2053-2051are not part of the main fatty acid catabolism pathway in *S. maltophilia*.

## DISCUSSION

Bacteria are commonly found in complex and heterogeneous communities. Within these environments, the exchange of signals governs the interactions and behaviors of the microbial populations. Typically, these functions involve both competitive and cooperative behaviors, such as production of virulence factors and intra- and interspecies communication (3, 42). The QS system in *S. maltophilia* is based on the fatty acid DSF, but it can also respond to exogenous AHLs (26). However, the effect of exogenous QS signals has not been studied in detail in this species. To this end, we supplemented early log-phase cultures of the model strain K279a with either DSF or AHLs at physiological concentrations and subjected each condition to RNA-Seq. Between 50% and 65% of the DEGs after treatment with each of the autoinducers, and just in the mid/late-exponential phase (6 h, OD_600_=1.5), are found also deregulated in non-induced cells collected in the stationary phase. These include genes required for energy generation pathways and stress response under oxygen-limiting conditions, β-oxidation, and iron storage, among others. It has been shown in other bacteria that QS regulates the transition into stationary phase in tight connection with starvation sensing (43), preparing the cells for a critical situation of stress due to lack of nutrients (44). The simple comparison of stationary-phase vs logarithmic-phase transcriptomes done in parallel, highlights the stressing environment of a high-density culture of *S. maltophilia*, which implies the activation of multiple stress pathways due to nutrient limitation and accumulation of metabolites, as seen in other bacteria (43, 45).

Among the most enriched COG categories found in both DSF and AHL conditions, those related with energy conversion are overrepresented, particularly those involving lipid and amino acid metabolism. This result could indicate that QS signals are primarily metabolized by *S. maltophilia*, and probably used as an energy source in complex and diverse communities. Remarkably, an overlap of genes whose expression was regulated by both DSF and AHLs was observed. These findings agree with previous studies in the unrelated *B. cepacia* complex and *B. cenocepacia* species, where the BDSF and AHL stimuli overlap through modulation of the intracellular levels of c-di-GMP (46, 47). QS systems in both these species control, among others, genes related with iron metabolism or proteins that use iron as a cofactor, as well as regulators of the stress response and general metabolism. Likewise, genes that fall in the same categories were found to be differentially expressed in the conditions here presented. The low amount of DEGs after induction with DSF in this study deserves to be discussed, considering that this autoinducer is sensed by the main QS system in *S. maltophilia*. The highest DSF production in these bacteria corresponds with a high-density culture just at the entrance of the stationary phase; however, trace amounts of DSF could be produced during early and mid/late stages of the exponential phase, which would help to explain the lower number of DEGs in the DSF condition as found in this study. In *Xcc* DSF production reaches a peak in the early-stationary phase, and the level subsequently declines, however transcription of *rpfF* is detected throughout the growth curve, with a marked increase in the entry of the stationary phase (15). Also, addition of DSF during exponential growth did not affect the subsequent expression of *rpf* genes in *Xcc*, indicating that DSF does not autoinduce its own synthesis (15). In our experiments, supplementing the cultures with DSF also did not impact the regulation of the *rpf* cluster, at least in the *rpfF*-1 variant strain K279a.

Among the DEGs that were found upregulated in all tested conditions, we selected the transcriptional regulator of the TetR/AcrR family (Smlt2053) for functional characterization. EMSA and transcriptomic analysis showed that Smlt2053 can recognize a consensus DNA sequence found in its own regulatory region and regulate the expression of its own operon through long-chain fatty acid sensing. Basically, Smlt2053 binds to its target regulated genes repressing their expression at low intracellular fatty acid concentration. The best characterized ortholog in other species is PsrA from *P. aeruginosa*. In this species, PsrA was found to recognize long chain fatty acids, through the C-terminus, and derepress its own expression upon the presence of the ligand (48, 49). Moreover, PsrA activates RpoS, the stationary phase sigma factor and it also regulates the expression of LexA (38, 40). Contrary to our results in *S. maltophilia*, transcriptome analysis of the *ΔpsrA* mutant in *P. aeruginosa* showed a broader network of genes whose expression is modulated by PsrA (41). However, that work suggested the involvement of PsrA in the regulation of β-oxidative enzymes, as shown here for Smlt2053.

The closest ortholog of Smlt2053 in the related species *Xanthomonas citri*, the TetR regulator XAC2014, was found to be the target of TfmR (T3SS and Fatty acid Mechanism Regulator), also supporting its role in the fatty acid pathways and virulence (36). Smlt2053 and XAC2014 harbor a palindromic DNA binding motif in the upstream promoter region, which is almost identical to the one characterized in *P. aeruginosa* (G/CAAAC(N_2-4_)GTTTG/C). All this, together with the fact that they both form an operon with two genes coding for enzyme variants of the β-oxidation pathway, suggests a similar role in lipid catabolism in the *Xanthomonadales* family. Interestingly, DSF showed the same effect as long chain fatty acids as an inducer of repressor Smlt2053. This result suggests that Smlt2053 could act as an intracellular receptor of DSF, acting as a key regulator of the signal turnover, by leading the excess of FAs to the β-oxidation pathway. An enzymatic breakdown of DSF within the cell has already been suggested by others after noticing that DSF concentration declines as the stationary phase progresses (15).

Although Smlt2053 is a repressor of its own operon, RNA-seq data shows a set of downregulated genes in the *Δsmlt2053* mutant which indicates that Smlt2053 plays a role in the activation of other genes. This finding could suggest a dual function as repressor and activator, or a secondary metabolite from the activity of enzymes Smlt2052 and Smlt2051 could modulate the expression of these genes. A dual function that also seems to have PsrA in *P. aeruginosa* (41). Among the few genes found to be downregulated in the mutant are the operon Smlt1443-Smlt1445 and the gene coding for the oxygen-binding protein Bacteriohemerythrin. The operon carries genes whose orthologs code for a transporter that is involved in the export of virulence factors in other species (50). Interestingly, the Bacteriohemerythrin gene was found upregulated in the stationary phase and after the induction with both DSF and AHLs. The ortholog of *S. maltophilia* Bacteriohemerythrin in *P. aeruginosa* plays a role in growth under microoxic conditions, particularly in a mutant of the QS regulator LasR (51). Adaptation of bacterial cells to the microoxic environment is important at the stationary phase where oxygen concentration dissolved in the growth medium is low due to the high cell density.

This work, in addition to giving clues about the regulation mechanism of QS in *S. maltophilia*, paves the way to find molecules that inhibit QS-dependent virulence factors. The QS system seems to confer a fitness advantage in high-density growth conditions, such as biofilm formation, and this could be modulated through quorum quenching (QQ). In particular, *S. maltophilia* isolates have shown QQ activity against AHLs. There is no report of DSF QQ activity in *S. maltophilia*, although it has been observed in other species (23, 26, 52). Fatty acids are now recognized as a potential alternative to conventional antibiotics because they have shown antibiofilm and antivirulence activity (53). It is known that one of the main mechanisms of action of candidate lipidic molecules in the cell is the QQ, therefore a therapeutic intervention that modulates lipid-mediated QS pathways in *S. maltophilia* could be an unconventional alternative to currently available antimicrobials.

## MATERIALS AND METHODS

### Bacterial strains and reagents

All bacterial strains used in this study are listed in supplementary Table S1. *S. maltophilia* K279a (54) was used as the reference strain in this study. K279a was routinely grown at 37°C in lysogeny broth (LB) medium on a rotatory shaker at 200 rpm, unless otherwise stated. Synthetic QS signals (N-octanoyl-HSL, C8-HSL; N-(3-Oxooctanoyl)-HSL, 3OC8-HSL; N-decanoyl-HSL, C10; and cis-11-methyl-2-dodecenoic acid, DSF) were obtained from Sigma-Aldrich. Stocks were prepared in DMSO at a concentration of 20 mg/ml, aliquoted and stored at -20°C until use. Free fatty acids used in this study were purchased from Sigma-Aldrich in the sodium salt form. Sodium butyrate and sodium octanoate were dissolved in water; sodium dodecanoate and 13-methyltetradecanoic acid (iso-C_15:0_) were dissolved in water containing 1% Brij-58; and sodium palmitate was dissolved in 10% pre-warmed methanol.

### Culture conditions and RNA extractions for RNA-seq

Two separate RNA-seq experiments were performed: 1) comparison of the changes in global gene expression in *S. maltophilia* K279a under QS-inducing conditions, and 2) transcriptome analysis of K279a mutant strain with a deletion of the Smlt2053 gene (K279aΔ*smlt2053*). All experiments were performed with triplicate samples. For the first experiment, *S. maltophilia* K279a was grown overnight at 37°C in LB to prepare fresh 150 mL LB cultures with an initial OD_550_ of 0.05. Cultures were grown to an OD_550_ of 0.2 (∼1.5 hours, beginning of the exponential phase) and supplemented with the corresponding QS autoinducers, as follows: AHL condition consisted of a combo of C8-HSL, 3OC8-HSL and C10-HSL, at 10 µM each; DSF was added at a final concentration of 10 µM; and the remaining flasks were supplemented with the same volume of DMSO to be used as the control. Cultures were then grown until they reached an OD_550_ of 1.5 (∼ 3 additional hours, transitioning from exponential to stationary growth phase) and harvested for RNA isolation. For the comparison of exponential phase (LOG) and stationary phase (STAT), RNA from triplicate cultures grown in LB were extracted at OD_550_ of 1.5 and 5, respectively. As for the comparison of K279a WT vs K279aΔ*smlt2053*, starting cultures and extraction time were identical to those of the previous experiment (collected at OD_550_ of 1.5). In all experiments total RNA was purified with the RNeasy Mini Kit (Qiagen), following the manufacturer’s instructions. The quantity and quality of each RNA sample were assessed with the 2100 Bioanalyzer (Agilent Technologies). Only samples with an RNA integrity number (RIN) > 8 were accepted and selected for library construction.

### RNA library construction and sequencing

The enrichment of bacterial mRNA was performed by depletion of the bacterial rRNA from total RNA extracted from *S. maltophilia* cultures. The rRNA was removed using the Ribo-Zero Bacteria Kit (Illumina) or MICROBExpress Bacterial mRNA Enrichment Kit (Thermo Fisher Scientific) starting with 500ng or 100ng, respectively, of bacterial total RNA. The RNASeq libraries were prepared following the TruSeq Stranded mRNA Library Prep (Illumina) protocol modified by discarding the polyA enrichment step. In brief, the enriched bacterial mRNA fraction was fragmented by divalent metal cations at elevated temperature. In order to achieve the library directionality, the second strand cDNA synthesis was performed in the presence of dUTP. The blunt-ended double stranded cDNA was 3’
sadenylated and Illumina platform compatible adaptors with indexes were ligated. The ligation product was enriched with 15 PCR cycles and the final library was validated on an Agilent 2100 Bioanalyzer (Agilent Technologies) using the DNA 7500 kit (Agilent Technologies).

The libraries were sequenced on HiSeq 2500 (Illumina) or HiSeq 4000 (Illumina) in paired-end mode with a read length of 2×76bp, using TruSeq SBS Kit v4 (Illumina) or HiSeq 4000 SBS kit (Illumina) respectively and following the manufacturer’s protocol. Images analysis, base calling and quality scoring of the run were processed using the manufacturer’s software Real Time Analysis (RTA 1.18.66.3 or RTA 2.7.7, respectively), followed by generation of FASTQ sequence files.

### RNA-seq data and sequence analysis

FastQC was used to verify the read quality of the RNA-Seq libraries and reads were mapped against the K279a genome using the STAR aligner. Gene abundances were normalized by calculating Fragments Per Kilobase Million (FPKM). The differential gene expression analysis was performed using the DESeq2 R package using an adjusted p-value < 0.05 (FDR of 5%) as a cut-off for statistical significance, with the Benjamini-Hochberg correction for multiple testing (55). To remove unwanted variation, the SVA method (56) was used. For subsequent analyses, *locus* with fold-change ≥ 2.0 were assigned as differentially expressed genes (DEGs). The raw, de-multiplexed reads as well as coverage files have been deposited in the NCBI’s Gene Expression Omnibus under the project ID: GSE206442 and GSE206554.

*S. maltophilia* K279a proteome was functionally annotated using the eggNOG-mapper (http://eggnogdb.embl.de/#/app/emapper) to assign each protein to a Cluster of Orthologous Groups (COG) category (57). COG categories for the K279a genome were obtained via the IMG (Integrated Microbial Genome) database (https://img.jgi.doe.gov/). The enrichment analyses consisted in a pathway and functional annotation of all DEGs with the TopGO R package (v2.38.1) and the KOBAS software (v3.0), for the GO and KEGG analysis, respectively (58, 59). Motif identification from the list of DEGs was performed by using the MEME suite (60). For a maximum of 50 top upregulated or downregulated genes in each comparison, the region spanning from -250bp to +50bp of the annotated ATG start codon was extracted and submitted to MEME analysis. Enriched motifs were then compared to known transcription factor binding sites with TOMTOM, whereas the K279a genome was used to scan for these motifs in the PRODORIC database (http://www.prodoric.de). Inverted repeats in nucleotide sequences were identified with the Palindrome program from the EMBOSS suite (https://www.bioinformatics.nl/cgi-bin/emboss/palindrome).

### Quantitative real-time PCR (qPCR) analysis

Gene expression and analysis was performed to determine the expression ratios of *smlt0264, smlt2944* and *smlt2052* in *S. maltophilia* K279a wild-type and mutated strain Δ*smlt2053* using specific primers (Table S1). Total RNA was isolated in three different experiments under the same conditions as for the RNA-seq experiments. One microgram of RNA was used to synthesize the cDNA with the Maxima Reverse Transcriptase kit (Thermo Scientific). qRT-PCR was performed with the CFX96 real-time PCR system (Bio-Rad). PCR products of ca. 150 bp were amplified for each gene and *rpoD* was used as a housekeeping control to normalize the gene expression levels. Expression was determined by the 2^-ΔΔCT^ method (61).

### Construction of unmarked deletion mutants

Markerless *S. maltophilia* K279a mutants were constructed according to the method initially developed for multidrug-resistant *B. cenocepacia*, using the pGPI-SceI/pDAI-SceI-SacB system (62). Briefly, this method is based on the I-SceI homing endonuclease system, which relies on two independent crossover events to generate markerless deletion mutants. First, the suicide plasmid pGPI-SceI-XCm (54, 63) containing a I-SceI recognition site and the flanking regions of the target region to be deleted (see supplementary Table S1 for primer details and plasmid construction) is introduced into the wild-type strain of *S. maltophilia* by triparental mating. After this first crossover event, the second plasmid pDAI-SceI-SacB is introduced. This second plasmid expresses the I-SceI endonuclease, which resolves the co-integrate structure with a second recombination event, stimulating the DNA repair machinery of the host strain to produce the markerless mutants. The pDAI-SceI-SacB plasmid is cured by sucrose counter-selection. Primers external to the deleted region (supplementary Table S1) were used to verify the mutants by both PCR and Sanger sequencing.

### Protein production and purification

To obtain the purified protein Smlt2053 the entire coding sequence was cloned, along with its upstream promoter region, into expression plasmid pET28a-TEV (primers listed in supplementary Table S1) which was electroporated into *E. coli* BL21(DE) as the expression host systems. IMAC purification was used to obtain the recombinant protein. Briefly, the resulting pellet of a 2L overnight culture was resuspended in 5 mM imidazole, 300 mM Nacl, 20 mM Tris-HCl, pH 7.9. After three rounds of sonication the sample was centrifuged for 45 minutes at 15,000g to separate the soluble and insoluble fractions. The soluble fraction was purified in a His-Trap (GE) column prior to the gradient elution in an FPLC ÄKTA pure platform. All the expression and purification procedures were performed by the Protein Production Platform (PPP) from Nanbiosis facilities at the BB-MRB from the UAB. Protein identity was confirmed by peptide mass fingerprinting (MALDI-TOF) at the Proteomics Laboratory from the CSIC/UAB, Barcelona (Spain).

### Electromobility shift assays (EMSA)

EMSA was adapted from the protocol of Hellman and Fried (64). DNA fragments containing the promoter regions of the target genes were generated by PCR using specific primers (Table S1). Binding reactions were carried out in a Tris-Glycine buffer (10 mM-40 mM) containing 20 mM KCl, 7.5% Glycerol, 0.01 mg/ml BSA and 1 mM DTT. Reaction mixtures consist of 50 ng of the DNA probe, 0 to 300 ng of purified protein Smlt2053 and 20 ng/µL poly (dI·dC). When required, different length fatty acids were included in the binding reaction at varying concentrations that ranged from 1 µM to 250 µM. The same solvent used to dissolve each fatty acid was used as a control in each assay. When the reaction mixture included a testing compound, the DNA-protein binding was incubated for 20 min before addition of the compound.

Reaction mixtures were incubated 30 min at 4°C and separated by electrophoresis on a native 6% polyacrylamide gel (Acrylamide/Bis-Acrylamide 37.5:1) in 0.5X TBE buffer (pH 8.3) containing 2.5% glycerol at 4°C in a cold room. After electrophoresis, gels were rinsed with water and stained with a 0.5X TBE solution containing 1X RedSafe (Sigma-Aldrich) for 15 minutes. To remove the exceeding dye, gels were washed twice with water for 10 minutes. Gels were visualized in a VersaDoc imaging system (Bio-Rad).

### Analysis of cellular fatty acids

Cellular fatty acids analysis was done at the Spanish Type Culture Collection (CECT, University of Valencia, Spain). Cells were grown on LB medium for 24 h at 37°C; extractions and determinations were performed according to the standard protocol of the MIDI Microbial Identification System (MIS; Microbial ID Inc., Newark, USA) (65) using an Agilent 6850 gas chromatograph (Agilent Technologies) following the Sherlock TSBA6 method and library.

## Data Availability

A full list of significant DEGs from each experimental condition is provided as a data supplement, and all raw sequence data are available at the NCBI’s Gene Expression Omnibus under the accession numbers GSE206442 and GSE206554.

## ACKNOWLEDGEMENTS

This work was funded by the Spanish MICINN (PID2019-111364RB-I00). Authors also thank Catalan AGAUR (2017 SGR 1062). Research at CNAG-CRG was funded by the Instituto de Salud Carlos III Spain (reference PT17/0009/0019) and co-funded by FEDER funds (European Regional Development Fund).

## Author contributions

X.C, P.H, M.B, O.C-S, A-C.G, A.E-C, M.D and M.G conducted the experiments. X.C, P.H, X.D, D.Y e I.G designed the experiments and participated in the analysis and interpretation of experimental data. M.D processed all the RNA-seq data and A.E-C performed the RNA-seq analysis. X.C, P.H, and D.Y wrote the paper. X.D, D.Y e I.G supervised research, and all authors revised the manuscript.

